# The molecular architecture of CenH3-deficient holocentromeres in Lepidoptera is dependent on transcriptional and chromatin dynamics

**DOI:** 10.1101/2020.07.09.193375

**Authors:** Aruni P. Senaratne, Héloïse Muller, Kelsey A. Fryer, Ines A. Drinnenberg

**Affiliations:** Institut Curie, PSL Research University, CNRS, UMR3664, F-75005 Paris, France; Sorbonne Université, Institut Curie, CNRS, UMR3664, F-75005 Paris, France; Department of Biochemistry, Stanford University School of Medicine, 279 Campus Drive, Beckman Center 409, Stanford, CA 94305-5307; Department of Genetics, Stanford University School of Medicine, Stanford, CA 94305-5120

## Abstract

Despite their essentiality for chromosome segregation, centromeres are diverse among eukaryotes and embody two main configurations: mono- and holocentromeres, referring respectively to a localized or unrestricted distribution of centromeric activity. Previous studies revealed that holocentricity in many insects coincides with the loss of the otherwise essential centromere component CenH3 (CENP-A), suggesting a molecular link between the two events. In this study, we leveraged recently-identified centromere components to map and characterize the centromeres of Bombyx mori. This uncovered a robust correlation between centromere profiles and regions of low chromatin dynamics. Transcriptional perturbation experiments showed that low chromatin activity is crucial for centromere formation in *B. mori*. Our study points to a novel mechanism of centromere formation that occurs in a manner recessive to the chromosome-wide chromatin landscape. Based on similar profiles in additional Lepidoptera, we propose an evolutionarily conserved mechanism that underlies the establishment of holocentromeres through loss of centromere specificity.

## Introduction

The centromere of each chromosome in a eukaryotic cell precisely defines the point of spindle fiber attachment in order to accurately partition the genome during each cell division. In the first chronological descriptions of cell division dating back to 1882, Walther Flemming observed that upon condensation, mitotic salamander chromosomes displayed a single region of narrowed chromatin to which spindle fibers physically attached. He termed this region as the primary constriction of a chromosome, thus representing a single (mono) site of centromeric activity (Flemming, 1882). In 1888, Theodor Boveri described that the mitotic chromosomes of the roundworm, *Ascaris megacephala* behaved differently to what had been described earlier by Flemming. *Ascaris* chromosomes lacked a primary constriction and had spindle fibers attached chromosome-wide, thus representing an unrestricted distribution of centromeric activity (Boveri, 1888). These two observations are the hallmarks of the two most common centromere architectures that we know of today, which are described as monocentric and holocentric, respectively.

Most eukaryotes harbor monocentric chromosomes. Monocentromeres range from the simplest ~125 base pair point centromeres of budding yeast (Carbon and Clarke, 1984) to the regional centromeres found abundantly in mammals and plants that can span up to several megabases in size (McKinley and Cheeseman, 2016; Muller et al., 2019). Holocentric organisms are also considerably widespread and found in diverse eukaryotic lineages such as nematodes, arthropods, and flowering plants (Melters et al., 2012).

The selective forces favoring either a mono- or holocentric architecture of chromosomes are unclear. However, the nested occurrence of holocentricity within larger monocentric groups in the eukaryotic tree has supported a general consensus that holocentric chromosomes in extant organisms were derived from monocentric ancestors (Escudero et al., 2016; Melters et al., 2012). The lack of conclusive evidence pointing to reversions to the monocentric form in any eukaryotic lineage further supports the unidirectional evolution of holocentricity and suggests it to be a stable trait once adapted. To explain the evolutionary transition from mono- to holocentric chromosomes, various, often species-specific, models and evolutionary drivers have been proposed (Malik, 2002; Nagaki et al., 2005; Neumann et al., 2012). Nevertheless, the underlying mechanism remains an open question.

Studies in a few select organisms have provided details into the molecular architecture of holocentric chromosomes. Such studies relied on profiling the distribution of the widely conserved centromere-specific histone H3 variant, CenH3 (first identified as CENP-A in mammals) (Earnshaw and Rothfield, 1985; Palmer et al., 1991) and have revealed that holocentromere organization can be different on the molecular level despite their common centromeric configuration. The elaborately described holocentric model nematode *Caenorhabditis elegans* has been shown to have low occupancy CenH3 domains spanning transcriptionally-silent chromatin interspersed among discrete, high occupancy CenH3 sites (Gassmann et al., 2012; Steiner and Henikoff, 2014). In contrast, in some holocentric plants, families of satellites and/or retrotransposons (Haizel et al., 2005; Marques et al., 2015) are found to partially co-localize with centromeres, although a centromere-specific role for these repeat sequences is unknown.

The presence of holocentric chromosomes has also been reported in several lineages of insects (Melters et al., 2012; Drinnenberg et al., 2014). Here, in contrast to holocentric nematodes and plants, protein homology predictions across mono- and holocentric insect orders revealed that multiple, independently derived holocentric insects do not possess CenH3 (Drinnenberg et al., 2014). This finding was unexpected given that the CenH3 protein is essential for chromosome segregation in most other eukaryotes. Together with its deposition factor HJURP/Scm3 (Dunleavy et al., 2009; Foltz et al., 2009) (CAL1 in flies (Chen et al., 2014)) and other kinetochore components, CenH3 takes part in a self-propagating epigenetic loop enabling the replenishment of its replication-dependent dilution at centromeres over cell divisions (Karpen and Allshire, 1997). With the exception of budding yeast and its close relatives where centromere location is defined genetically (Carbon and Clarke, 1984, Dujon et al., 2004; Kobayashi et al., 2015), the participation of CenH3 in this epigenetic loop has been demonstrated in several monocentric systems including human cells (Barnhart et al., 2011; Black et al., 2004; Carroll et al., 2010; Fachinetti et al., 2013; Guse et al., 2011; Kato et al., 2013; Logsdon et al., 2015; Tachiwana et al., 2015) and flies (Mendiburo et al., 2011; Palladino et al., 2020; Roure et al., 2019). Alternatively, a genetic basis of centromere identity has also been proposed through the recognition of tertiary DNA structures by the CenH3 incorporation machinery, with the incorporation of CenH3 itself being a by-product of this process (Kasinathan and Henikoff, 2018). Regardless of an epigenetic or genetic basis of centromere definition, the centromere-specific function of CenH3 is found to be essential in all organisms tested (Blower and Karpen, 2001; Buchwitz et al., 1999; Howman et al., 2000; Stoler et al., 1995; Talbert et al., 2002).

In addition to multiple lineages of holocentric insects, cases of CenH3 loss have also been reported in trypanosomes (Akiyoshi and Gull, 2014; Talbert et al., 2009) and in an early-diverging fungus (Hooff et al., 2017; Navarro-Mendoza et al., 2019). Interestingly, the multiple CenH3 gene loss events in several orders of insects that are notably concomitant with the occurrence of a holocentric architecture of chromosomes in each case is the first common event to be reported surrounding the loss of CenH3. This co-occurrence of holocentromeres with CenH3 loss suggests that these insects employ alternative, CenH3-independent modes of centromere specification that occur genome-wide. Given their monocentric ancestry, CenH3-deficient insects therefore represent a useful experimental tool to understand the molecular mechanism that underlies the transition to holocentric architectures in these organisms.

In this study, we aimed to determine how CenH3-deficient holocentromeres are defined in a representative insect order, Lepidoptera (butterflies and moths). We approached this question using a combination of imaging, genomics and chromatin perturbation analyses to map and characterize the centromeres in the silk moth, *Bombyx mori*. Our findings revealed a link between centromere location and regions of low chromatin activity along the entire length of *B. mori* chromosomes. Based on our results, we propose a model for how the loss of centromere specificity can lead to chromosome-wide establishment of centromeres that is non-randomly defined by underlying chromatin activity states. Our study provides insights into the conditions governing the loss of centromere specifying components such as CenH3 over evolutionary timescales.

## Results

### The kinetochore forms a broad localization pattern along *B. mori* mitotic chromosomes

To understand holocentromere architecture in CenH3-deficient insects, we used a cell line derived from our representative insect model system, *B. mori*. We targeted one previously identified centromere-proximal component of the *B. mori* kinetochore, CENP-T (Cortes-Silva et al., 2020) to visualize its localization pattern on mitotic *B. mori* chromosome spreads by immunofluorescence (IF) microscopy using a custom-made antibody (Cortes-Silva et al., 2020). We observed that the CENP-T-specific IF signal formed broad localization patterns along the polar length of sister chromatids (Figure 1A), a pattern reminiscent to kinetochore staining in other holocentric organisms (Buchwitz et al., 1999). While CENP-T is an inner kinetochore component that binds directly to DNA as shown in vertebrates (Hori et al., 2008), the centromere-distal outer kinetochore components bridge centromeric DNA to the spindle apparatus, thus serving as a proxy for potential sites of spindle fiber attachment (Musacchio and Desai, 2017). To test whether the broad localization pattern we observed for CENP-T corresponds also to sites of outer kinetochore assembly, we co-stained our chromosome spreads with another custom-made antibody specific for the outer kinetochore component Dsn1 (Cortes-Silva et al., 2020). We found that the CENP-T and Dsn1 immunosignals co-localize with one another (Figure 1B), demonstrating that diffuse regions along the poleward surface of *B. mori* sister chromatids represent potential sites driving chromosome segregation during mitosis. These stainings are the first visualizations of kinetochore components along *B. mori* chromosomes, thereby building on previous cytological data (Murakami and Imai, 1974) and confirming that *B. mori* chromosomes are holocentric.

**Figure 1:**
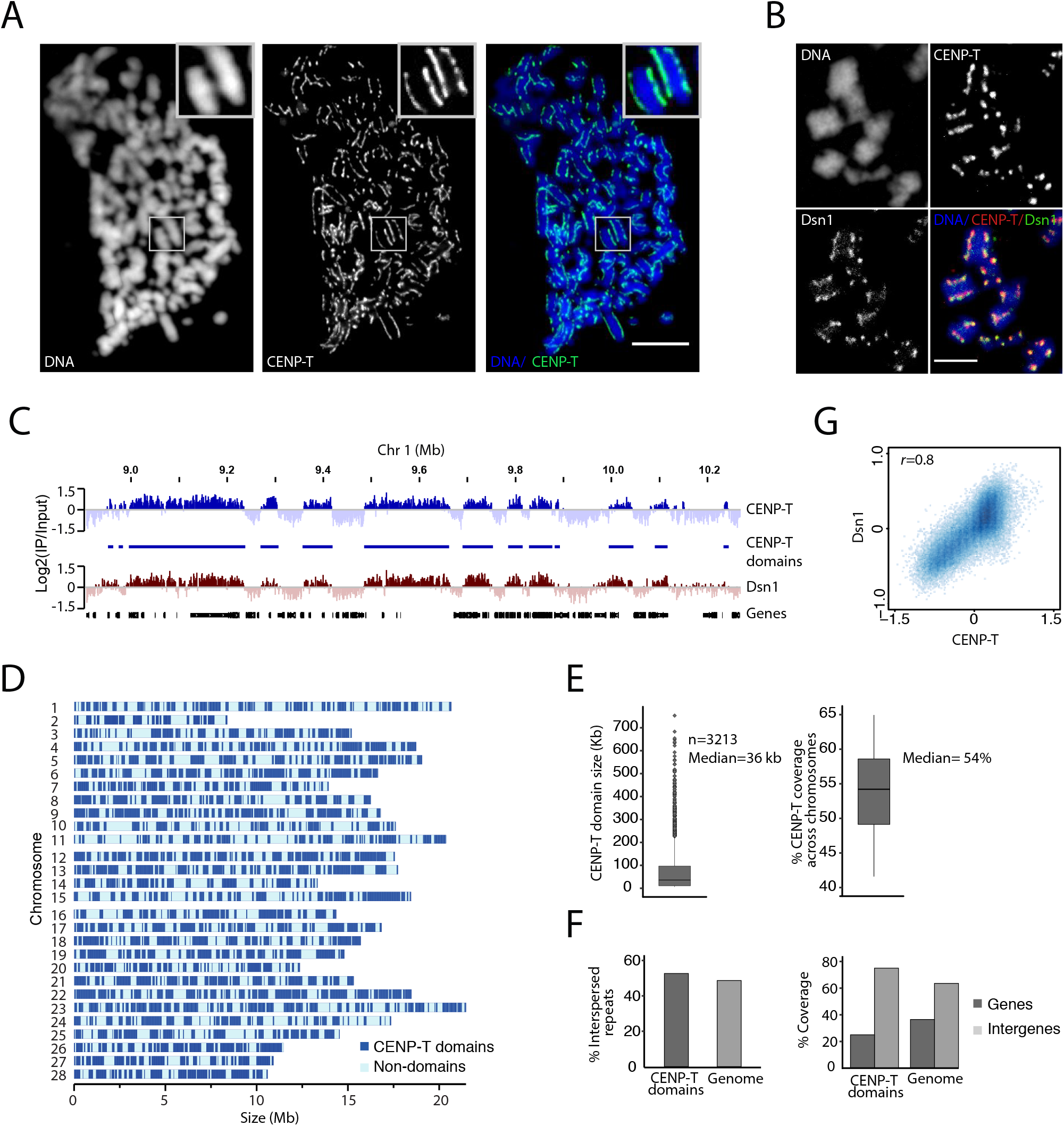
Kinetochore localization patterns in *B. mori*. **A)** Representative IF image of *B. mori* DAPI-stained mitotic chromosomes (blue) showing broadly-distributed immunosignal patterns of CENP-T (green). Scale bar: 5μm **B)** Representative IF image of *B. mori* DAPI-stained mitotic chromosomes (blue) showing the immunosignal patterns of CENP-T (red) and Dsn1 (green). Co-localization of CENP-T and Dsn1 signals can be distinguished as yellow foci in the overlay. Scale bar: 5 μm. **C)** Genome-browser snapshot of a representative portion of *B.mori* chromosome 1 for CENP-T X-ChIP-seq, CENP-T domains, Dsn1 X-ChIP-seq and annotated genes. ChIP-seq signals are represented as histograms of the average log2 ratio of IP/Input in genome-wide 1 kb windows. **D)** Size-scaled schematics of 28 *B. mori* chromosomes showing the distribution of CENP-T domains (dark blue segments). **E)** Features of CENP-T domains: boxplots showing the sizes (left) and the genome coverage in percent by CENP-T domains (right). **F)** Genomic features underlying CENP-T domains: barplots showing the fraction in percent of annotated interspersed repeats within CENP-T domains (dark grey) and genome-wide (light grey) (left); and the fraction in percent of annotated genes within CENP-T domains (dark grey) and genome-wide (light grey) (right). **G)** Genome-wide correlation plot of CENP-T and Dsn1 occupancy. Average log2 ratios of IP/Input in 10 kb windows were used for plotting and calculating the pearson correlation coefficient (*r*) indicated on the top-left corner.

### Half of the *B. mori* genome is permissive for kinetochore assembly

We proceeded to perform chromatin immunoprecipitation followed by sequencing (ChIP-seq) experiments targeting CENP-T in unsynchronized *B. mori* cell populations in order to obtain genomic resolution maps of CENP-T’s distribution on chromatin. These analyses revealed broad regions of CENP-T enrichment (Figure 1C) with maximum enrichment scores of about 2-fold (log2) over input, indicating an extensive, yet low level of CENP-T localization along chromosomes. This pattern was highly reproducible across replicates (Figure S1A and S1B). Given that the ChIP-seq libraries were prepared from asynchronous cell populations containing ~ 2% mitotic cells (Cortes-Silva et al., 2020) and that the *B. mori* CENP-T, as the vertebrate CENP-T homologs (Hori et al., 2008) also localizes to chromatin during interphase (Figure S1E), CENP-T interphasic localization likely contributes to the majority of its enrichment signal.

To evaluate the extent to which CENP-T-enriched sites identified by ChIP-seq are functional kinetochore assembly sites to drive chromosome segregation during mitosis, we performed ChIP-seq to map the genome-wide distribution pattern of Dsn1. As other outer kinetochore components, Dsn1 is found on centromeric chromatin only during mitosis (Musacchio and Desai, 2017). Analogous to our co-stainings of CENP-T and Dsn1 on mitotic chromosome spreads, the ChIP-seq profile of Dsn1 also correlated well with that of CENP-T (*r* = 0.8) (Figure 1C, G).

To better characterize CENP-T-binding patterns, we used a custom domain-calling pipeline to annotate CENP-T domains. This method revealed that CENP-T domains show no obvious clustering towards the center or telomeric regions of chromosomes (Figure 1D); are of variable size (median size = 36 kb); and have a genome-wide median coverage of 54% (Figure 1E). Considering that CENP-T coverage is quantified for CENP-T ChIP-seq signal from a cell population, this number reflects the average CENP-T coverage for that population. Thus, while approximately half of the genome is CENP-T-permissive, it is possible that from one cell to the next, CENP-T occupies only a fraction of permissive sites.

Next, we analyzed the distribution of repeat sequences underlying CENP-T sites. Approximately 47% of the *B. mori* genome is comprised of repetitive DNA sequences, with around 97% of these repeats being Class I and Class II transposable elements (Kawamoto et al., 2019). To determine if CENP-T domains preferentially occur in these repetitive elements, we first intersected our annotated CENP-T domain coordinates with a database of consensus transposon sequences for *B. mori* (kind gift from the Pillai Lab, University of Geneva). This approach revealed that the proportion of transposon sequences underlying CENP-T domains is not significantly different to that in the rest of the genome (Figure 1F). Second, as a complementary approach, we searched *de novo* for any centromere-enriched repeats as described (Smith et al., 2020). Consistent with the first approach, these analyses also revealed that CENP-T-permissive sites in the *B. mori* genome are not enriched for repetitive DNA sequences (Figure S1F). Additionally, intersecting the CENP-T domain coordinates with *B. mori* gene annotations (obtained from SilkBase: http://silkbase.ab.a.u-tokyo.ac.jp) revealed that CENP-T domains are relatively depleted in genes (Figure 1F). Collectively, these results led us to conclude that *B. mori* centromeres are organized as broad domains that are composed of complex DNA and depleted from gene bodies.

### Kinetochore attachment sites in *B. mori* are anti-correlated with actively transcribed chromatin

Given the broad distribution pattern and absence of any consensus sequence underlying CENP-T sites, it is unlikely that centromeres along *B. mori* chromosomes are defined by a specific DNA sequence. Our use of genomics approaches to profile the underlying chromatin environment instead revealed several correlations of the distribution of CENP-T with respect to chromatin marks governing transcriptional status. We found a positive correlation between the distributions of CENP-T and tri-methylated Lysine 27 on histone H3 (H3K27me3) (*r* = 0.6) (Figure 2A, B), a histone mark that is typically associated with transcriptionally-silent heterochromatin (Kouzarides, 2007). Conversely, we found a negative correlation in the distributions of CENP-T and two histone marks that are associated with a transcriptionally-active chromatin state (Kouzarides, 2007): (i) tri-methylated Lysine 4 on histone H3 (H3K4me3) (*r* = −0.2), and (ii) tri-methylated Lysine 36 on histone H3 (H3K36me3) (*r* = −0.6) (Figure 2A, B). We also profiled the distribution of di-and tri-methylated Lysine 9 on histone H3 (H3K9me2 and H3K9me3, respectively), two histone marks that are typically associated with transcriptionally silent heterochromatin (Kouzarides, 2007). However, the resulting ChIP-seq profiles generated using multiple different antibodies and ChIP protocols for H3K9me2/3 were very similar to a histone H3 ChIP-seq profile (*r* = 0.8 for H3K9me2 *vs* H3; and *r* = 0.7 for H3K9me3 *vs* H3, respectively) (Figure S2B, C). This offered us limited confidence in the observed patterns of H3K9me2/3 in our cell line. However, as opposed to the H3K9me3-enriched heterochromatin blocks found at the pericentromeres of monocentric organisms (Sullivan and Karpen, 2004), a lack of such regions was observed in our *B. mori* cell line as evident in the absence of any chromocenters in interphase nuclei (Figure S2D).

**Figure 2:**
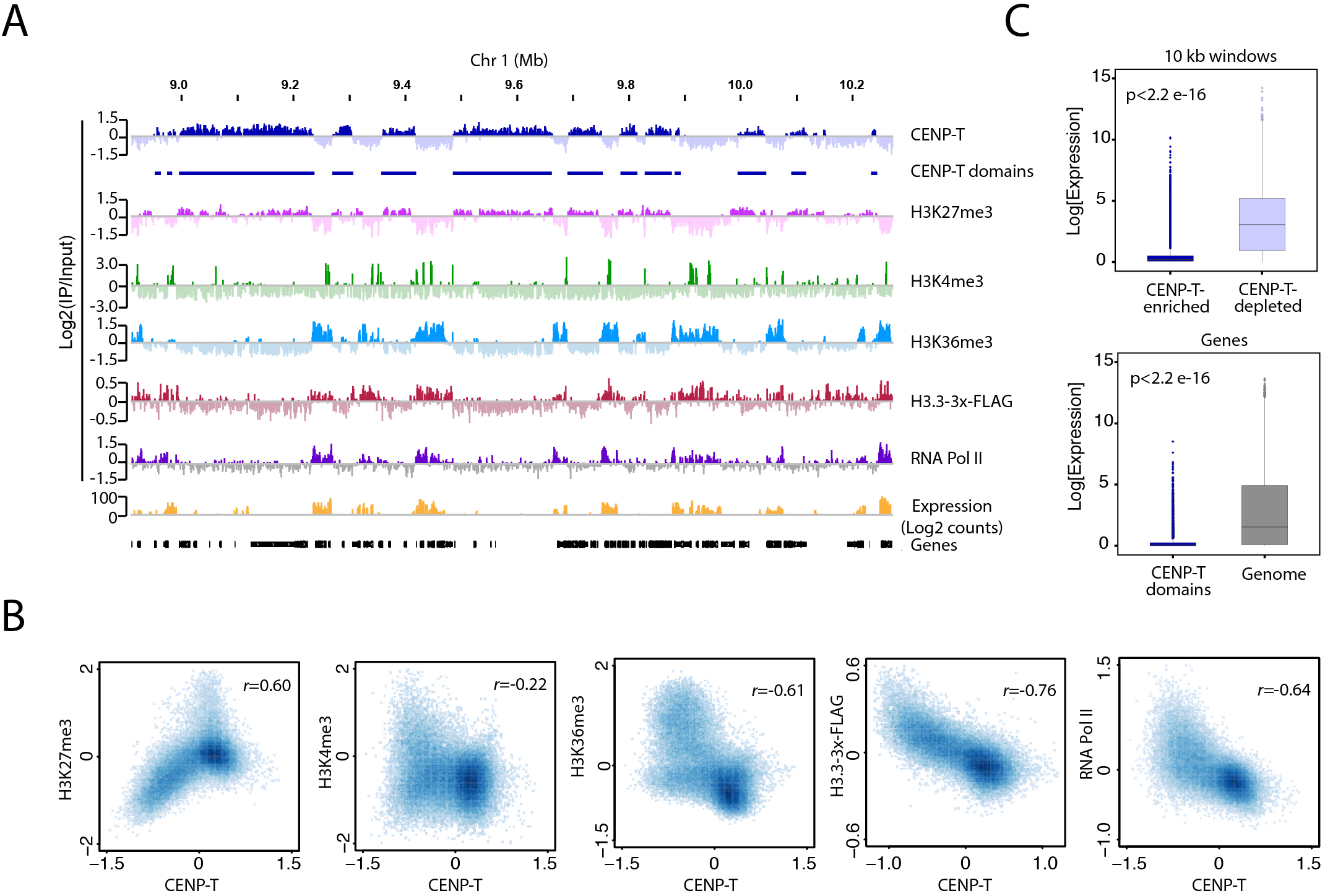
*B. mori* kinetochore attachment sites are anti-correlated with actively transcribed chromatin and nucleosome turnover. **A)** Genome-browser snapshot of a representative portion of *B. mori* chromosome 1 for CENP-T X-ChIP-seq, CENP-T domains, H3K27me3 N-ChIP-seq, H3K4me3 X-ChIP-seq, H3K36me3 N-ChIP-seq, H3.3-3X-FLAG N-ChIP-seq, RNA Pol II X-ChIp-seq, mRNA-seq and annotated genes. ChIP-seq signals are represented as histograms of the average log2 ratio of IP/Input in genome-wide 1 kb windows. mRNAseq signal is represented as a histogram of log2 normalized counts per million mapped bins (BPM). **B)** Genome-wide correlation plots comparing the occupancy of CENP-T and H3K27me3, H3K4me3, H3K36me3, H3.3-3X-FLAG and RNA Pol II. Average log2 ratios of IP/Input in 10 kb windows were used for plotting and calculating the pearson correlation coefficient (*r*) indicated on the top-right corner of each plot. Comparisons between N-ChIP-seq and X-ChIP-seq or X-ChIP-seq replicates for the histone marks are shown in Figure S2A. **C)** Top: boxplot showing the expression levels in genome-wide 10 kb windows that are CENP-T-enriched (dark blue) or CENP-T-depleted (light purple), and bottom: boxplot showing the expression levels in annotated genes that are 100% within CENP-T domains (dark blue) as compared to annotated genes genome-wide (grey). Expression levels are represented as the log2 normalized transcripts per million mapped reads (TPM). Statistical significance was tested using the Kolmogorov-Smirnov test.

We additionally generated mRNA-seq data for our cell line in order to assess the expression levels at *B. mori* kinetochore attachment sites. We found that mapped transcripts negatively correlated with CENP-T domains (Figure 2A). Additionally, annotated genes that did fall completely within CENP-T domains had significantly lower expression levels (median = 0.06 normalized expression units) compared to the genome-wide average (median = 1.53 normalized expression units) (Figure 2C). To account for transcripts mapping to non-annotated genes, we also looked at the expression levels across the entire genome. As with the gene-level analysis, we found that genomic regions that are enriched for CENP-T have significantly lower expression values (median = 0.2 normalized expression units) than CENP-T-depleted regions (median = 3.06 normalized expression units) (Figure 2C). Consistent with our ChIP-seq profiles for histone marks and our mRNA-seq profiles, the ChIP-seq profile of RNA Polymerase II (RNA Pol II) also revealed a negative correlation with CENP-T throughout the genome (*r* = −0.64) (Figure 2A, B). Finally, we also profiled epitope-tagged histone H3.3 driven under a constitutive promoter as a measure of nucleosome turnover (Kraushaar et al., 2013). Compared to our RNA Pol II and active histone mark profiles, we found that the H3.3 profile shows the strongest anti-correlation with CENP-T (*r* = −0.76) (Figure 2A, B). Thus, our results indicate that centromeres are excluded from genomic regions undergoing active nucleosome turnover, driven by transcription or other chromatin remodeling processes (Figure 2A, B).

### Hormone–induced perturbations of gene expression result in proximal CENP-T loss or gain

To test our hypothesis that there is a causal link between *B. mori* centromere location and chromatin activity, we used an approach that allowed us to study the effects of induced transcriptional changes on CENP-T localization patterns. In insects, the ecdysone hormone response is a well-defined transcriptional response leading to the systematic activation or repression of a subset of specific genes involved in metamorphoses and development (Yamanaka et al., 2013).

We treated asynchronous *B. mori* cells with 20-Hydroxyecdysone (20E) for 48 hours and carried out mRNA-seq to determine whether our cell line had an effective transcriptional response (Figure 3A). Indeed, we found that while the expression pattern in annotated genes remained overall well-correlated before and after 20E treatment (r = 0.99), several genes showed differential expression upon treatment (as indicated by clusters forming away from the diagonal in the +/− 20E correlation scatterplot; Figure 3B). To identify a subset of differentially expressed genes for further analyses, we applied cut-offs in expression level to define up- and down-regulated genes (See Methods). Among a subset of twenty-six up-regulated genes, we found known 20E-responsive genes including orphan nuclear receptor genes (Yamanaka et al., 2013) (Figure 3B). We also identified one down-regulated gene that corresponded to a 20E-hydroxylase (Figure 3B). The functional annotations of these genes furthered our confidence in the specificity of the 20E response in our cell line.

**Figure 3:**
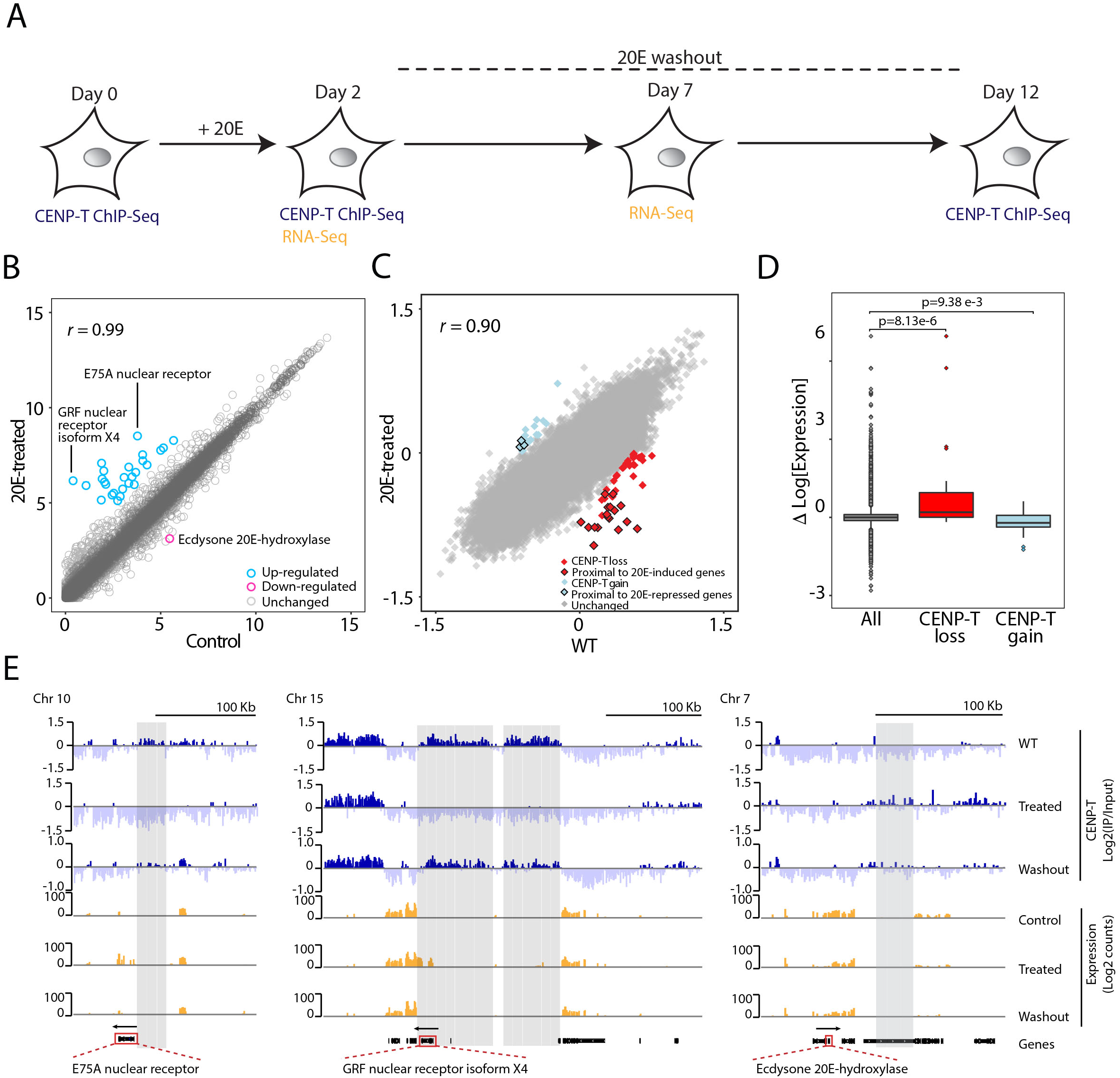
Hormone–induced perturbations of gene expression result in proximal CENP-T loss or gain. **A)** Schematic summarizing the steps of 20E treatment and 20E washout in BmN4 cells. **B)** Genome-wide correlation plot comparing the expression levels of annotated genes (grey circles) in 20E-treated vs control (DMSO-treated) conditions. The subset of twenty-six differentially-expressed genes with user-defined expression cut-offs is highlighted (pink and blue circles, respectively). Functions of two up-regulated genes and one down-regulated gene that could be linked to three cases of differential CENP-T occupancy in the 20E-treated condition are annotated in the plot. Log2-transformed transcripts per million mapped reads (TPM) are used to represent expression level per gene and to calculate the pearson correlation coefficient (*r*) indicated on the top-left corner. **C)** Genome-wide correlation plot comparing CENP-T occupancy before (WT) and after 20E treatment in 10 kb windows (grey boxes). 10 kb windows with differential CENP-T occupancy with user defined enrichment cut-offs are demarcated (red or blue boxes, respectively). Average log2 ratio of IP/Input in 10 kb windows were used for plotting and for calculating the pearson correlation coefficient (*r*) indicated on the top-left corner. Only those 10 kb windows with total loss or total gain of CENP-T after 20E treatment from a previously enriched or depleted state, respectively were considered for our analyses. **D)** Boxplot: difference in expression levels between 20E treatment and control in genome-wide 10 kb windows (grey) and the subset of 10 kb windows with depleted or enriched CENP-T occupancy (red and blue, respectively). Difference in expression was calculated by subtracting the log2 TPM scores of control from 20E-treated for each 10 kb window. Statistical significance was tested using the Kolmogorov-Smirnov test. **E)** Genome-browser snapshots of *B. mori* chromosomes showing two cases of CENP-T loss (left & middle) and one case of CENP-T gain (right) that could be linked to proximal changes in gene expression. Tracks are shown for CENP-T X-ChIP-seq profiles (blue) for WT, 20E-treated and 20E-washout conditions and RNA-seq profiles (orange) for control, 20E-treated and 20E-washout conditions. Pre-identified 10 kb windows with differential CENP-T occupancy (grey boxes) were found to fall consecutively within these regions. The three differentially-expressed 20E-specific genes lying upstream of these regions are highlighted with a red box in the gene track. Gene direction is marked with a black arrowhead. ChIP-seq signal is represented as the log2 ratio of IP/Input in genome-wide 1 kb windows. RNA-seq signal is represented as log2 normalized counts per million mapped bins (BPM). Genome-browser snapshots showing CENP-T occupancy around the remaining twenty-three pre-identified differentially-expressed genes are shown in Figure S3D.

Having confirmation of a visible change in expression, we next profiled CENP-T occupancy under the same conditions (Figure 3A). To identify any changes in CENP-T localization, we compared CENP-T enrichment patterns across genome-wide 10 kb windows between treated and untreated conditions. Consistent with the largely unaltered transcriptome in annotated genes (Figure 3B) and 10 kb windows (Figure 3D), global CENP-T occupancy levels were also well-correlated in treated and untreated conditions (*r* = 0,9) (Figure 3C). Nevertheless, we were able to identify specific loci with altered CENP-T levels after treatment (Figure 3C). Comparing mRNA expression levels in genomic regions showing lost or gained CENP-T binding after 20E treatment revealed that CENP-T depleted regions showed elevated expression while CENP-T enriched regions showed decreased levels of expression (Figure 3D). This is consistent with our above findings that CENP-T binding is more robust in transcriptionally silent regions.

Next, we zoomed in on each of the individual loci that lost or gained CENP-T after treatment in order to further interpret each CENP-T loss or gain event with respect to the broader chromosomal environment. We found that the most pronounced cases of CENP-T loss (representing 17 out of 48 CENP-T-depleted 10 kb genomic windows) were in fact in the proximity of two up-regulated genes. These loci corresponded to large consecutive regions of CENP-T loss after 20E treatment on chromosomes 10 and 15 where in each case, the losses were just upstream of the genes encoding for a 20E-specific nuclear receptor protein (Figure 3E). On the other hand, when we zoomed in on a genomic region on chromosome 7 that gained CENP-T after 20E treatment, we observed the opposite scenario, where the gain in CENP-T was in close proximity to a down-regulated 20E-hydroxylase gene (Figure 3E). While we cannot explain all of the observed changes in CENP-T occupancy after 20E treatment, the above described examples of CENP-T gain or loss near genes with altered expression allowed us to partially link CENP-T localization patterns to changes in transcriptional activity. The fact that altered CENP-T occupancy did not necessarily co-localize with the altered transcriptional output, but rather extended to large upstream regions, reinforces our hypothesis that changes of the chromatin landscape such as nucleosome eviction or reassembly in gene bodies, promoter or enhancer regions of up- or down-regulated genes interfere with or enable CENP-T localization, respectively.

In order to further evaluate whether it is a change in chromatin dynamics in the proximity of differential-regulated genes that underlies the changes in CENP-T occupancy, we allowed the transcriptional program to reset over the course of 5 days upon removal of the 20E hormone from the media (Figure 3A). Analyses of gene expression levels in the hormone-washout sample revealed that the previously identified subset of induced genes was mostly restored to expression levels similar to that of the control gene subset (Figure S3A). To evaluate the effects of restored gene expression on CENP-T localization, we once again profiled CENP-T occupancy 10 days following hormone-washout (Figure 3A). We found that upon 20E-washout, the prominent CENP-T losses on chromosomes 10 and 15 were recovered to levels similar to wild-type profiles (Figure 3E). The recovered events of previous CENP-T loss extended upstream of the now repressed mRNA output, once again indicating a link between chromatin activity/status and CENP-T deposition. The CENP-T gain upon 20E-treatment that was proximal to the down-regulated 20E-hydroxylase gene on chromosome 7 did not completely recover, which could be explained by incomplete restoration of expression of this gene (Figure 3E). In addition to these individual cases, similar levels of recovery following 20E-washout were observed genome-wide for regions with differential CENP-T occupancy (Figure S3B) along with comparable expression levels across all genomic windows (Figure S3C).

Taken together, our results support the hypothesis that the dynamics of the chromatin landscape determined by promoter activation or Pol II passage govern CENP-T occupancy such that CENP-T is removed from active chromatin. The complete loss of CENP-T upon promoter and gene activation also shows that no immediate recycling mechanism exists to restore CENP-T levels in those regions. Finally, the recovery of CENP-T localization upon 20E washout indicates that restoring low chromatin dynamics is sufficient for centromere formation.

### Differentially expressed orthologous genes in Lepidoptera show opposite patterns of CENP-T localization

Given that the same chromosomal locus could be made permissive or repressive to CENP-T by merely changing the underlying chromatin activity status, we reasoned that such opposite patterns of CENP-T enrichment should be readily observable in an experimentally unperturbed setting, for example, on a gene that naturally varies in expression levels. To address this possibility, we turned to a close lepidopteran relative of *B. mori*, the cabbage looper *Trichoplusia ni*. We hypothesized that differentially-expressed orthologous genes between these two species provide a natural template of varying expression levels at orthologous loci to which we can correlate CENP-T occupancy levels (Figure 4A).

**Figure 4:**
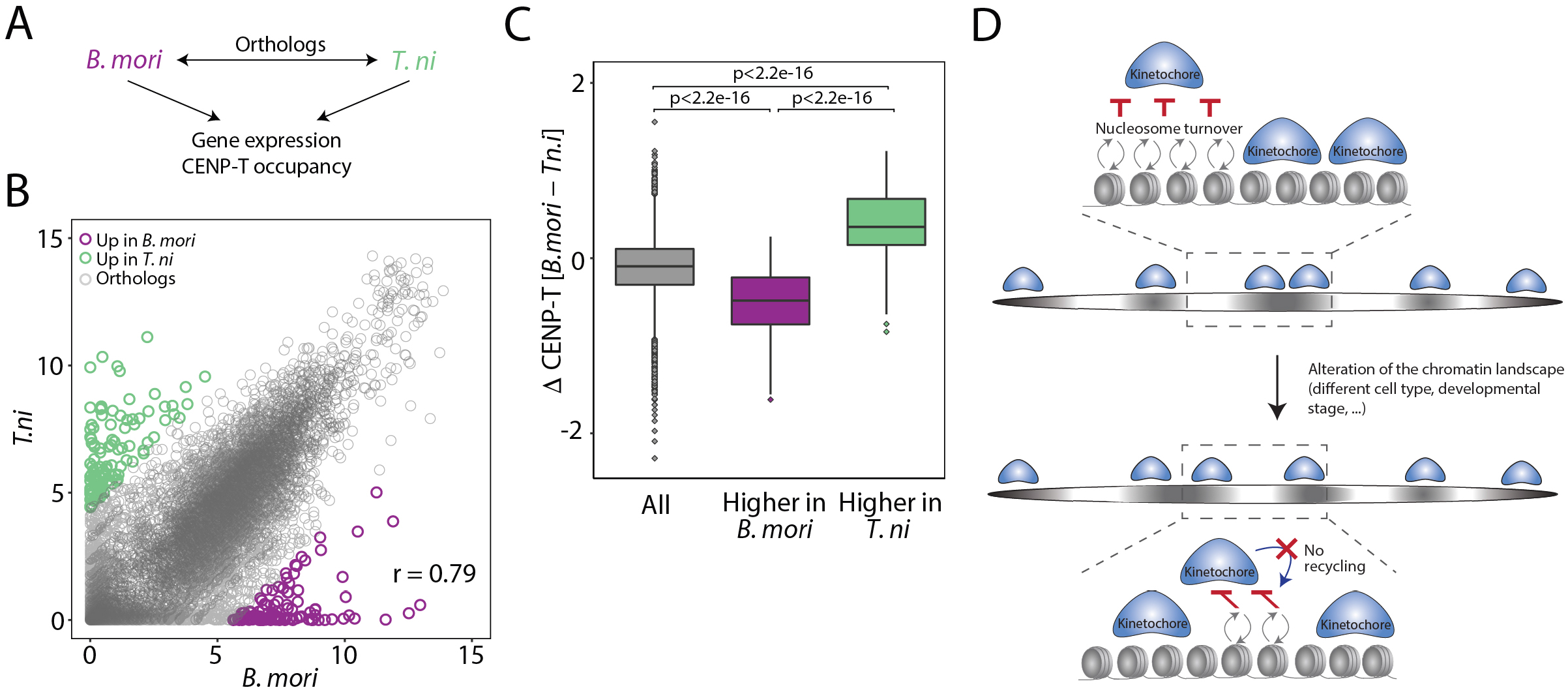
Differentially expressed orthologous genes in Lepidoptera show opposite patterns of CENP-T localization. **A)** Schematic summarizing the concept to identify a link between CENP-T occupancy and transcriptional activity by using naturally differentially-expressed orthologous genes of *B. mori* and *T. ni* as a template. Correlation plot showing the expression level of orthologous genes in *B. mori* and *T. ni*. Genes with similar expression levels (gray circles) and those corresponding to the subsets of the most highly expressed genes that were identified in each species which are silent in the corresponding species (purple or green circles) are indicated. Expression level per gene is represented as the log2-transformed number of transcripts per million mapped reads (TPM) score and was used to calculate the pearson correlation coefficient (*r*) indicated on the bottom-right corner. **C)** Boxplot: difference in CENP-T occupancy across all orthologs (gray) and across the subset of highly expressed genes of *B. mori* and *T. ni* (purple and green, respectively). Difference in CENP-T occupancy per gene was calculated by subtracting the average log2 IP/Input scores for CENP-T ChIP-seq in *B. mori* from that of *T. ni*. Statistical significance was tested using the Kolmogorov-Smirnov test. **D)** Model for the architecture of CenH3-deficient holocentromeres in insects. Kinetochores bind chromosome-wide due to lack of centromere specificity. Binding is opposed only by high nucleosome turnover. Any changes to the chromatin landscape that result in alterations to the nucleosome turnover profile will also alter kinetochore attachments toward more stable regions. This kinetochore binding profile is not epigenetically inherited from one cell cycle to the next, therefore resulting in different kinetochore binding patterns for every different nucleosome turnover profile.

To first evaluate whether centromeres in *T. ni* are defined in a similar way to *B. mori*, we profiled CENP-T in an unsynchronized germ cell line derived from *T. ni* (Granados et al., 1986). We found that the distribution of *T. ni* CENP-T ChIP-seq signal resembled that of *B. mori,* wherein CENP-T localized to broad chromosomal regions (Figure S4A). Next, we did ChIP-seq profiling of the histone marks H3K27me3 and H3K36me3 in our *T. ni* cell line, which we used as markers of transcriptionally-repressed and -active chromatin, respectively. Similar to *B. mori,* in *T. ni*, the genome-wide distribution of CENP-T was positively and negatively correlated with H3K27me3 (*r* = 0.5) and H3K36me3 (*r* = −0.5), respectively (Figure S4A, B) and showed a negative correlation to mapped mRNA transcripts (Figure S4A). Thus, we concluded that the CENP-T localization in *T.ni* is correlated with silent chromatin similar to what is seen in *B. mori*.

We then reciprocally searched the *B. mori* and *T. ni* proteomes with one another to select a set of encoding orthologous genes. This revealed > 9500 orthologs to which we applied a fold-change and cut-offs in expression to select a subset of differentially-expressed genes for each species (See Methods). Accordingly, we selected 115 genes that are expressed in *B. mori* (and low in *T. ni*) and 110 genes that are expressed in *T. ni* (and low in *B. mori)* (Figure 4B). We then quantified the average CENP-T ChIP-seq scores for *B. mori* and *T. ni* over both sets of genes. In line with our previous observations, we found that CENP-T ChIP-seq signal was depleted over those genes that are expressed in *B. mori,* while CENP-T ChIP-seq signal was enriched over the corresponding lowly expressed orthologs of *T. ni* (Figure 4C). In a similar manner, we observed that CENP-T ChIP-seq signal was enriched over those genes that are lowly expressed in *B. mori,* while CENP-T ChIP-seq signal was depleted over the corresponding expressed orthologs of *T.ni*. We have thus demonstrated that across orthologous genes, CENP-T occupancy is significantly different only in those cases where there is differential expression, further supporting our hypothesis linking centromere formation to the underlying chromatin activity status.

## Discussion

In this study, we characterized the CenH3-deficient holocentromere architecture of Lepidoptera, which allowed us to make an important link between their chromatin landscape and centromere distribution. Our use of a perturbation system to show that this landscape is critical for creating centromere-permissive or -dismissive environments along lepidopteran chromosomes provides insights into a new mode of centromere definition independent of CenH3.

More precisely, we identified recurrent patterns of negative correlations between the genome-wide distribution of CENP-T and factors associated with high nucleosome turnover including RNA Pol II and H3.3. We propose that lepidopteran centromere formation does not depend on an active recruitment process involving a centromere-specific epigenetic or genetic factor. Instead, kinetochores can assemble non-specifically and anywhere along the chromosomes where nucleosome turnover is low. Changes in gene expression and thus the chromatin landscape can thereafter disrupt kinetochore attachment leading to its complete loss due to the absence of an active recycling mechanism. The CENP-T profile that we thus measure corresponds to attached kinetochores that could persist over the cell cycle. This is different to the dynamics of H3.3 during transcription for instance, where a combination between new H3.3 deposition and recycling of pre-existing H3.3 coordinated by the HIRA complex enables the maintenance of the epigenetic information at that locus (Torné et al., 2019). In contrast, the presence of the lepidopteran kinetochore is not memorized and can change from cell cycle to cell cycle dependent on the chromosome-wide chromatin landscape (Figure 4D).

The organization and inheritance of the Lepidopteran holocentromere might be conceptually similar to the CenH3-encoding holocentromere in *C. elegans*. It has been proposed that actively transcribed chromosomal domains are refractory to CenH3 incorporation in *C. elegans* late-stage embryos likely due to elevated nucleosome turnover in those regions (Gassmann et al., 2012; Steiner and Henikoff, 2014). Instead, CenH3 nucleosomes can stably remain over chromosomal domains with low levels of RNA Pol II occupancy. Given the observation that *C. elegans* CenH3 was completely turned-over at each cell cycle and discontinued in the germline, the authors further proposed that CenH3 might not propagate centromere identity in this organism (Gassmann et al., 2012). The ubiquitous negative relationship between active chromatin and centromeres throughout the genomes of CenH3-deficient Lepidoptera closely resembles that of *C. elegans*. That is, based on our chromatin-perturbation experiments, we can conclude that the induction of transcriptional activity or chromatin remodeling is sufficient to induce complete dissociation of CENP-T (Figure 3). In turn, CENP-T can accumulate *de novo* in regions without requiring pre-existing CENP-T as cues (Figure 3). The potential similarities in centromere regulation in the absence or presence of CenH3 highlight the relevance of chromatin dynamics for holocentromere organization across species.

### Evolutionary establishment of chromosomes with holocentric architecture in insects

The findings of this study allow us to propose a molecular mechanism underlying the mono-to holocentric transition in Lepidoptera and possibly other holocentric insects. Ancestral insects were monocentric and CenH3-dependent (Drinnenberg et al., 2014; Melters et al., 2012). CenH3 in ancestral monocentric insects might have self-propagated centromere identity at a restricted locus where it nucleated kinetochore assembly, resembling centromeres in *Drosophila melanogaster* and other monocentric organisms (Barnhart et al., 2011; Black et al., 2004; Carroll et al., 2010; Fachinetti et al., 2013; Guse et al., 2011; Karpen and Allshire, 1997; Kato et al., 2013; Logsdon et al., 2015; Mendiburo et al., 2011; Palladino et al., 2020; Roure et al., 2019; Tachiwana et al., 2015). Alternatively, centromeres could have been defined by a genetic mechanism (Kasinathan and Henikoff, 2018). In contrast, in the CenH3-deficient derived state that we characterize in this study, kinetochore assembly and thus, centromere activity occurs chromosome-wide, only antagonized by chromatin disruption processes (Figure 4D). Given that CenH3 in insects is essential in monocentric Diptera (Blower and Karpen, 2001) and its loss is only found in holocentric lineages (Drinnenberg et al., 2014), the establishment of this derived holocentric state is likely to have preceded and subsequently allowed the loss of CenH3. This is also supported by the fact that some holocentric Hemipteran insects have CenH3 homologs (Cortes-Silva et al., 2020). These Hemipterans could therefore represent an intermediate form leading to the establishment of the CenH3-deficient state.

While the molecular function of CenH3 during this possible transition state is an open question, two events must have occurred to allow the progression to a CenH3-deficient state: (i) centromere identity either conferred through an epigenetic feedback loop or through a genetic definition of centromere identity was lost, which led to the establishment of a holocentric architecture; (ii) kinetochore assembly on DNA must have become CenH3-independent, perhaps through the replacement of its ability to attach the kinetochore complex to chromatin by other kinetochore components. Future studies aiming to characterize the centromere architecture of additional holocentric insects will give more resolution to these intermediate events.

It is plausible that *C. elegans* and other holocentric nematodes that encode CenH3 may resemble this transitional stage of centromere organization. Compared to the complex CCAN of holocentric insects that mediates CenH3- and CENP-C-independent kinetochore assembly (Cortes-Silva et al., 2020), the inner kinetochores of nematodes lack CCAN but contain CenH3 and its direct binding partner CENP-C (Buchwitz et al., 1999; Cheeseman, 2004; Moore and Roth, 2001) explaining their critical roles for kinetochore attachment and outer kinetochore assembly (Carroll et al., 2010; Oegema et al., 2001; Cheeseman, 2004; Desai, 2003; Milks et al., 2009; Przewloka et al., 2011; Screpanti et al., 2011).

## Acknowledgements

We thank Andreas Rechtsteiner and Nicolas Servant for helpful discussions regarding the bioinformatic pipelines used in this study, Munetaka Kawamoto and Susumu Katsuma for early access to new functional annotations for the gene models of *B. mori*, Camille Berthelot for helpful discussions regarding the analyses of orthologs, Mickaël Garnier and Patricia Le Baccon for help with imaging by microscopy, the Almouzni lab for kindly gifting us with mouse ES cells, all members of the Fachinetti and Drinnenberg labs as well members of A.P.S’s thesis committee; Geneviève Almouzni and Leah Rosin for helpful discussions. APS receives salary support from Institut Curie and the Fondation Recherche Médicale. IAD receives salary support from the CNRS. This work is supported by the Labex DEEP ANR-11-LABX-0044 part of the IDEX Idex PSL ANR-10-IDEX-0001-02 PSL, an ATIP-AVENIR Research grant, Institut Curie and the ERC (CENEVO-758757).

