## SupplementalData for "The molecular architecture of CenH3-deficient holocentromeres in Lepidoptera is dependent on transcriptional and chromatin dynamics"

#### Supplemental figure legends

**Figure S1: CENP-T localizes to non-repetitive domains during interphase and mitosis.**

**A)** X-ChIP-seq profiles are well-correlated across varied MNase conditions. Genome-wide correlation plot of CENP-T occupancy from replicate X-ChIP-seq profiles generated with 2 Units of MNase and 45 mins of digestion. Average log2 ratios of IP/Input in genome-wide 10 kb windows were used for plotting and calculating the pearson correlation coefficient (*r*) indicated on the top-left corner. **B)** Genome-browser snapshot of a representative portion of *B. mori* chromosome 11 for CENP-T X-ChIP-seq at varied MNase conditions, FLAG X-ChIP-seq (negative control), and annotated genes. Time and concentration of MNase are annotated adjacent to each track. ChIP-seq signal is represented as the average log2 ratio of IP/Input in genome-wide 1 kb windows. **C)** Agarose gel image of nucleosome enrichment profiles for input and CENP-T pull-down at 15 mins, 30 mins, 45 mins, and 60 mins of MNase digestion (2 units), respectively. IPs from chromatin subject to 30 mins and 45mins of MNase digestion from the same experiment were sequenced and resulting X-ChIP-seq profiles are shown above in Figure S1B tracks 2 and 3, respectively. **D)** Validation of CENP-T antibody specificity. Representative images of BmN4-SID1 mitotic cells in either WT condition (top panel) or after three days of RNAi treatment targeting CENP-T (bottom panel). Cells were stained for DNA (left), pre-validated CENP-T antibody generated in rabbit host (middle) (see Cortes-Silva et al., 2020) and CENP-T antibody generated in mouse host (right) used in this study for co-stainings wth Dsn1. Scale bar: 5 μm. **E)** CENP-T is present in interphase. Quantifications of mean fluorescence intensity of CENP-T as compared to control cells stained for secondary Alexa Fluor 568 antibody) in BmN4-SID1 interphase cells (n=10 cells). Statistical significance was tested using the Wilcoxon-Mann-Whitney test. **F)** CENP-T domains are non-repetitive. k-mer plots showing the normalized k-mer counts (green circles) in CENP-T X-ChIP-seq vs Input datasets for k-mer lengths of 10 bp (left) and 25 bp (right), respectively. Black, blue, green and red diagonal lines depict increasingly stringent k-mer enrichment ratio cut-offs.

**Figure S2: Histone mark patterns in *B. mori*.**

**A)** Genome-wide correlation plots of H3K27me3 N-ChIP-seq *vs* H3K27me3 X-ChIP-seq (Left); replicates of H3K4me3 X-ChIP-seq (middle); and H3K36me3 N-ChIP-seq *vs* H3K36me3 X-ChIP-seq (right). Average log2 ratios of IP/Input in 10 kb windows were used for plotting and calculating the pearson correlation coefficient (*r*) indicated on the top-right corner of each plot. **B)** Genome browser snapshot of a representative portion of *B. mori* chromosome 5 for CENP-T X-ChIP-seq, H3K9me2 N-ChIP-seq, H3K9me2 X-ChIP-seq I and II, H3K9me3 N-ChIP-seq, H3K9me3 X-ChIP-seq I and II, H3 N-ChIP-seq and annotated genes. The antibody used for each H3K9 IP is annotated adjacent to the respective track. ChIP-seq signal is represented as the average log2 ratio of IP/Input in genome-wide 1 kb windows. **C)**

Genome-wide correlation plots of H3K9me2 N-ChIP-seq or H3K9me3 N-ChIP-seq *vs* H3 N-ChIP-seq (top) and *vs* CENP-T X-ChIP-seq (bottom). Average log2 ratios of IP/Input in 10kb windows were used for plotting and calculating the pearson correlation coefficient (*r*) indicated on the top-right corner of each plot. **D)** Representative IF microscopy images of *B. mori* and mouse interphase cells stained for H3K9me3 (red) and DNA (blue). The anti-H3K9me3 antibodies that were used are annotated to the right. Chromocenters constituting H3K9me3-enriched heterochromatin can be seen as intensely DAPI- and H3K9me3-stained nuclear foci in the mouse cells for both antibodies, whereas homogenous DAPI staining can be seen for *B. mori.* Scale bar: 10μm.

**Figure S3: 20E-washout restores transcription and CENP-T occupancy levels.**
**A)** Genome-wide correlation plot comparing the expression levels of annotated genes (grey circles) in 20E-washout *vs* DMSO-control conditions. Pre-identified subset of twenty-six differentially-expressed genes upon 20E treatment are marked as pink and blue circles, respectively. Functions of two up-regulated genes and one down-regulated that were linked to three cases of differential CENP-T occupancy in the 20E-treated condition are annotated in the plot. Log2-transformed number of transcripts per million mapped reads (TPM) is used to represent expression level per gene and to calculate the pearson correlation coefficient (*r*) indicated on the top-left corner. **B)** Genome-wide correlation plot comparing CENP-T occupancy in 20E-washout *vs* WT conditions in 10 kb windows (grey boxes). Pre-identified 10 kb windows with differential CENP-T occupancy upon 20E-treatment are marked as red or blue filled boxes, respectively. Average log2 ratios of IP/Input in 10 kb windows were used for plotting and calculating the pearson correlation coefficient (*r*) indicated on the top-right corner. **C)** Boxplot: difference in expression levels between 20E-washout and DMSO-control in genome-wide 10 kb windows (grey) and in the subset of pre-identified 10 kb windows with depleted or enriched CENP-T occupancy after 20E treatment (red and blue, respectively). Difference in expression was calculated for each 10 kb window by subtracting the log2 TPM score of control from 20E-washout. Statistical significance was tested using the Kolmogorov-Smirnov test. **D)** Genome browser snapshots of *B. mori* chromosomes for CENP-T X-ChIP-seq (blue) and RNA-seq (orange) surrounding twenty-three remaining up-regulated genes (marked with a red box in the gene annotation track) that were identified with user-defined expression cut-offs. CENP-T X-ChIP-seq tracks are shown for WT, 20E-treated and 20E-washout conditions. RNA-seq tracks are shown for DMSO control, 20E-treated and 20E-washout conditions. ChIP-seq signal is represented as the average log2 ratio of IP/Input in genome-wide 1 kb windows. RNA-seq signal is represented as log2 normalized counts per million mapped bins (BPM).

**Figure S4: Centromere specification is conserved among CenH3-lacking Lepidoptera.**
**A)** Genome-browser snapshot of a representative portion of *T. ni* chromosome 4 for CENP-T X-ChIP-seq, H3K27me3 N-ChIP-seq, H3K36me3 N-ChIP-seq and annotated genes. ChIP-seq signal is represented as the average log2 ratio of IP/Input in genome-wide 1 kb windows. CENP-T ChIP in *T. ni* was carried out using the same validated CENP-T antibody used for *B. mori* ChIPs. **B)** Genome-wide correlation plots of *T. ni* CENP-T occupancy and H3K27me3 (left); and H3K36me3 (right). Average log2 ratios of IP/Input in 10 kb windows were used for plotting and calculating the Pearson correlation coefficient (*r*) indicated on the top- left and top-right corner of each plot, respectively.

#### Methods

##### Lepidopteran cell lines and culture conditions

Cultured silkworm ovary-derived BmN4 (ATCC catalog # CRL-8910; RRID: CVCL_Z633), BmN4-SID1(RRID:CVCL_Z091) (Kobayashi et al., 2012) and *T. ni* Hi5 cell lines were maintained in Sf-900 II SFM medium (GIBCO catalog # 10902-088) supplemented with (BmN4, BmN4-SID1) or without (Hi5) 10% fetal bovine serum (Eurobio catalog # CVFSVF0001), antibiotic-antimycotic (GIBCO catalog # 15240-062) and 2mM L-glutamine (GIBCO catalog # 25030-024) at 27°C.

##### Validation of *B. mori* CENP-T mouse antibody specificity by RNAi

BmN4-SID1 cells were grown in 24-well plates for three days with or without 400 pg/μL dsRNA targeting CENP-T. At the end of three days, RNAi-treated or Wild-type (WT) cells were fixed with 100% ice-cold MeOH for 10 min at -20°C and then processed for immunofluorescence (IF) microscopy.

##### Preparation of mitotic chromosome spreads

BmN4 cells were grown in 6-well plates and collected by mild centrifugation (300 g) at room temperature. Supernatant containing growth medium was decanted and cell pellet was gently re-suspended in hypotonic buffer (80% water, 20% PBS) that was added drop-wise under mild vortex to the cell pellet. Cells were incubated in hypotonic buffer for 30 minutes at room temperature. Swollen cells in hypotonic buffer were then aliquoted into disposable funnels and cytospun onto coverslips using a Shandon cytospin 3 centrifuge. Chromosome spreads were un-mounted from cytospin and were fixed immediately with 4% PFA for 10min at room temperature and processed for IF microscopy.

##### Immunofluorescence

Cells fixed in either 100% ice-cold MeOH (stainings in BmN4-SID1 cells) or 4% PFA (mitotic chromosome spreads and H3K9me3 stainings in BmN4 cells and mouse ES cells) were permeabilized using 0.3% Triton x-100 in PBS. Cells were then blocked in 3% BSA-PBS. Primary antibody incubations were done in blocking buffer overnight at 4°C. The following primary antibodies were used at 1:1000 dilution: anti-CENP-T serum (rabbit or mouse polyclonal), anti-Dsn1 serum (rabbit polyclonal) generated by Covalab (Villeurbanne, FR) (Cortes-Silva et al., 2020), and H3K9me3 antibodies: rabbit polyclonal, Abcam, ab8898 and mouse monoclonal, MBL MABI0318. The next day, cells were washed three times with 0.3% Triton x-100 in PBS and were incubated for 1 hour at 4°C with secondary antibodies diluted to 1:1000 in blocking buffer. The following fluorescent-conjugated secondary antibodies were used: goat anti-rabbit IgG Alexa Fluor 488 (Thermo Fisher Scientific, catalog # A-11034, RRID AB_2576217), goat anti-rabbit IgG Alexa Fluor 568 (Thermo Fisher Scientific, A-11011, RRID AB_143157), and goat anti-mouse IgG Alexa Fluor 568 (Thermo Fisher Scientific, catalog # A-11004, RRID AB_2534072). Cells were washed three times with 0.3% Triton x-100 in PBS and counterstained with DAPI for 3min at room temperature (Sigma catalog # D9542) before washing again in 0.3% Triton x-100 in PBS and mounting samples in Vectashield Antifade Mounting Medium (Vector Laboratories catalog # H-1000; RRID:AB_2336789).

**Microscopy**

Images of chromosome spreads and H3K9me3 stainings in mitotic or interphase BmN4 or mouse ES cells were acquired on a LSM780 confocal microscope. Z stacks were acquired at 0.1-0.2 μm intervals using the 100X oil objective. Images of BmN4-SID1 interphase cells stained for CENP-T in WT or RNAi conditions were acquired on a Zeiss Axiovert Z1 light microscope. Z stacks were acquired at 0.2 μm intervals using the 100X oil objective.

Quantification of fluorescence intensity was performed using the Fiji software (Schindelin et al., 2012) on unprocessed TIFF images. IF signal in interphase BmN4-SID1 cells stained for either CENP-T or Alexa Fluor 568 were quantified. Nuclei of 10 cells were manually segmented using DAPI signal. The mean fluorescence intensity of each nucleus was then measured and corrected for background. For background correction, the average of mean intensities of three random circular regions of fixed size (10x10 pixels) placed outside the nuclear areas was determined and subtracted from the CENP-T or Alexa-Fluor-specific IF signal of each nucleus.

**Cross-linked ChIPs (X-ChIP) using in-house protocol**

X-ChIP was performed as previously described (Skene and Henikoff, 2015) with the following modifications. Two confluent T75 flasks (Thermofisher, catalog # 156499) of BmN4 cells (for CENP-T, Dsn1, RNA Pol II and FLAG ChIPs) or Hi5 cells (for CENP-T ChIP) were used. Cells were cross-linked in freshly prepared 1% MeOH-free formaldehyde (Thermofisher catalog # 28906) for 10 min at room temperature. Cross-linking was quenched by adding glycine to 125 mM for 2 min at room temperature. Cells were then washed in ice-cold PBS and incubated for 10 min with 150 µl ice-cold lysis buffer (1% SDS, 10 mM EDTA, 50 mM Tris-HCl pH 8.1) with cOmplete Protease Inhibitor Cocktail (Roche catalog # 11697498001). To the cell lysates, 1350 µl ChIP buffer (1% Triton X-100, 150 mM NaCl, 2 mM EDTA, 20 mM Tris-HCl pH 8.1,) with Protease inhibitor cocktail was added along with 4.5 µl CaCl2 1M (3 mM final) and then pre-warmed for 2 min at 37 °C. Nuclei were then treated with 1 or 2 units of MNase (Sigma catalog # N3755-500UN) for 15 min (CENP-T and FLAG ChIPs), 30 min (CENP-T ChIP), 45 min (CENP-T, Dsn1, RNA Pol II ChIPs) or 60 min (CENP-T ChIP), respectively at 37 °C. MNase reaction was stopped by adding a mix of 30 µl EDTA (0.5 M stock) and 60µl EGTA (0.5 M stock). Each MNase-treated nuclei sample was then sonicated using a Covaris E220 sonicator under the following parameters: 150 sec (WT CENP-T ChIPs in BmN4 cells) or 250 sec (Dsn1, RNA Pol II and 20E CENP-T ChIPs in BmN4 cells ; CENP-T ChIP in Hi5 cells), Duty 10%, Power 75 W, cycles/burst 200, 7 °C. Sonicated chromatin was centrifuged 3 min at 16000 g and clear supernatant containing the solubilized chromatin was saved either as input or for ChIP. Anti-CENP-T serum (rabbit polyclonal), anti-Dsn1 serum (rabbit polyclonal) or anti RNA Pol II antibody (rabbit polyclonal, Abcam, ab817) diluted in PBST (0.3% Triton X-100 or 0.02% tween-20) was incubated with Protein A dynabeads (Thermofisher, catalog # 10001D) for 10 min at room temperature to allow for pA-beads-antibody binding  Antibody-bound beads were washed with ChIP buffer and mixed with input chromatin. Alternatively, commercial beads-anti-FLAG M2 antibody (Sigma catalog # M8823; RRID:AB_2637089) was directly added to input chromatin for control FLAG X-ChIP. All samples were incubated overnight at 4 °C. Chromatin-bound beads were collected the next day on a magnetic rack and washed with the following ice-cold buffers: once with low-salt TSE 1 (0.1% SDS, 1% Triton X-100, 150 mM NaCl, 2 mM EDTA, 20 mM Tris-HCl pH 8.1,); four times with high-salt TSE II (0.1% SDS, 1% Triton X-100, 500 mM NaCl, 2 mM EDTA, 20 mM Tris-HCl pH 8.1); and three times with 1x TE. DNA was directly extracted from chromatin-beads or input by adding DNA extraction buffer (20 mM Tris-HCl pH 8.1, 10 mM EDTA, 5 mM EGTA, 300 mM NaCl, 1% SDS) and incubating at 37°C followed by reversing cross-links by addition of proteinase K (Qiagen, catalog # 19131) and incubation overnight at 65°C. DNA was isolated with Phenol:Chloroform extraction and precipitated with NaOAc and 100% EtOH in the presence of glycogen. DNA was finally re-suspended in 1x TE containing RNase (1µg/µl) and incubated for 15 min at 37 °C. Nucleosome profiles for Input and ChIP DNA were analyzed using a Agilent bioanalyzer or Agilent 4200 Tapestation with a DNA high sensitivity kit.

**Cross-linked ChIPs using commercial protocol**

ChIP and DNA extraction was performed as described in the Diagenode iDeal ChIP-seq kit for histones (Diagenode, catalog # C01010051/ C01010057) using two confluent T75 flasks of BmN4 cells. Chromatin was sheared using a Covaris E220 sonicator. Solubilized chromatin was incubated overnight at 4 °C with the following antibodies: H3K4me3 (positive control ChIP-seq grade antibody provided in Diagenode iDeal ChIP-seq kit); H3K9me2 (mouse monoclonal, MBL, MABI0317; and rabbit polyclonal, Diagenode, C15410060); H3K9me3 (mouse monoclonal, MBL, MABI0318; and rabbit polyclonal, Abcam, ab8898); H3K27me3 (Rabbit polyclonal, Cell Signaling Technology, C36B11); and H3K36me3 (rabbit polyclonal, Abcam, ab9050). Nucleosome profiles for Input and ChIP were analyzed using a Agilent bioanalyzer or Agilent 4200 Tapestation with a DNA high sensitivity kit.

**Native ChIPs**

ChIP and DNA extraction were performed as described (Orsi et al., 2015) using two confluent T75 flasks of BmN4 or Hi5 cells.  Solubilized chromatin was incubated overnight at 4 °C with the following antibodies: H3K27me3 (Rabbit polyclonal, Cell Signaling Technology, C36B11); H3K36me3 (rabbit polyclonal, Abcam, ab9050); H3K9me2 (rabbit polyclonal, Activ Motif, AM39753); H3K9me3 (rabbit polyclonal, Abcam, ab8898); H3 (Rabbit polyclonal, Abcam, ab1791). Nucleosome profiles for Input and ChIP were analyzed using a Agilent bioanalyzer or Agilent 4200 Tapestation with a DNA high sensitivity kit.

**Next-generation sequencing and ChIP-seq data analysis**

All steps of Illumina library preparation and sequencing were carried out at the Curie Institute's sequencing platform. Adapter trimmed, single-end Illumina reads of 100 bp length were mapped using Bowtie2 (Langmead and Salzberg, 2012) with default parameters to the *B. mori* genome assembly (Kawamoto et al., 2019) downloaded from Silkbase: http://silkbase.ab.a.u-tokyo.ac.jp, which was modified to extract only assembled chromosomes 1 to 28 or the *T. ni* genome (Fu et al., 2018) downloaded from the Cabbage Looper Database: https://cabbagelooper.org. After removal of duplicates using Picard tools (http://broadinstitute.github.io/picard/), Deeptools (Ramirez, 2016) bamCompare function (Ramírez et al., 2016) was used to generate ChIP-seq signal tracks represented as histograms of the average log2-ratio of RPKM-normalized read counts in IP over Input in genome-wide 1 kb windows that were visualized in IGV (Robinson, 2011). Deeptools multiBigWigSummary function was used to compute average log2-ratio of RPKM-normalized read count in IP over Input in genome-wide 10 kb windows for making scatterplots (Wickham H (2016), https://ggplot2.tidyverse.org) of the correlation between different ChIP-seq targets. Pearson correlation was calculated in RStudio (RStudio Team (2016). RStudio:

Integrated Development for R. RStudio, Inc., Boston, MA URL http://www.rstudio.com/) after filtering out those 10 kb windows with zero mapped reads in both IP and Input.

**Annotation of CENP-T domains**

CENP-T ChIP-seq signal originally in 1 kb windows was averaged over 2 kb. The averaged windows with positive scores (log2-ratio>0) within a genomic distance of 5 kb were merged using BEDTools (Quinlan and Hall, 2010). This cut-off for merging distance was determined so that the likelihood of finding consecutive 1 kb windows with positive CENP-T signal was significant at p<=0.05. Average CENP-T ChIP-seq signal was re-computed over the merged coordinates after removing any merged intervals of size <5kb. Any merged intervals with overall negative ChIP-seq scores were further removed. These were defined as positive domains. Negative domains were similarly annotated using a reciprocal approach where log2 scores <=0 were considered as negative scores. BEDTools (Quinlan and Hall, 2010) was used to subtract negative domains from positive domains to extract final CENP-T domains.

**Repeat analyses using RepeatMasker**

The *B. mori* genome assembly (chromosomes 1 to 28) and newly-annotated CENP-T domains were searched using RepeatMasker software version 4.08 and RMBlast version 2.10.0+ (http://www.repeatmasker.org) with a custom library of consensus transposon sequences for *B. mori* (kind gift from the Pillai Lab, University of Geneva). Simple repeats and low-complexity repeats accounting for <1% of the genome (Kawamoto et al., 2019) were omitted from the analyses. The percentage of interspersed repeats in CENP-T domains and genome-wide was calculated as the total number of base pairs covered by interspersed repeats as a fraction of the total genome size.

**Repeats analyses using k-mer clustering**

k-mer based repeat analysis pipeline from the Straight Lab (Smith et al., 2020): Single-end Illumina reads (adapter-trimmed, PCR duplicates removed) for CENP-T ChIP-seq and Input were used to generate k-mer databases for each dataset at k-mer lengths of 10 bp and 25 bp. The abundancedof each k-mer in both datasets was counted and normalized to the total number of basepairs in that dataset. K-mers found fewer than 10 times in either dataset were excluded from the analysis. Normalized k-mer counts in the CENP-T dataset (y-axis) were plotted as a function of normalized k-mer counts in Input (x-axis). Enrichment values for each k-mer were calculated as the ratio of normalized count in CENP-T dataset over the normalized count in Input dataset. Different enrichment cut-offs were defined according to the number of median absolute deviations away from the median enrichment ratio.

**Transcriptional profiling**

Total RNA was isolated from 1 confluent T75 flask of BmN4 or Hi5 cells using Trizol reagent (Invitrogen, catalog # 15596018) following the manufacturer's instructions. PolyA-selected RNA-seq libraries were prepared using the Illumina Truseq stranded mRNA protocol and sequenced at the Curie Institute's sequencing platform. Adapter-trimmed, paired-end reads of 100 bp length were mapped to the *B. mori* genome assembly (chromosomes 1 to 28) using STAR version 2.7 (Dobin et al., 2013). The number of mapped RNA-seq reads in annotated genes (downloaded from SilkBase: http://silkbase.ab.a.u-tokyo.ac.jp) or in genome-wide 10 kb windows were counted using ht-seq software (Anders et al., 2015). Read counts were transformed into TPM (transcripts per million mapped transcripts) scores to evaluate normalized expression levels. 1 kb resolution TPM-normalized RNA-seq coverage tracks were further generated using Deeptools BamCoverage function for visualization in IGV as histograms.

**20E treatment and washout**

20E-treated or DMSO-control samples were prepared by growing BmN4 cells in 5 ug/ml of 20E-hydroxyecdysone (Sigma-Aldrich, H5142) or <1% DMSO for 48 hours, following which RNA-seq (20E-treated and DMSO-treated) or CENP-T X-ChiP-seq (20E-treated) was performed. For 20E-washout RNA-seq and ChIP-seq experiments, cell layers growing for two days in the presence of 20E were softly rinsed with fresh growth medium thrice to dilute out the hormone without disrupting the monolayer. Cell layers were then allowed to grow in conditioned growth medium* for a further 5 days (RNA-seq) or 10 days (ChIP-seq) before harvesting. Cells growing in conditioned medium were expanded as appropriate by splitting with 50% fresh growth medium + 50% conditioned medium.

* Conditioned medium was collected from WT BmN4 cells growing for the same time period as the 20E treatment (2 days). Medium was decanted into a falcon tube and centrifuged at 1000 g for 10 min to remove any cells in the medium. Supernatant was filtered twice through a 0.22 μm filter and stored at 4°C for a maximum of 3 hours until use.

**Criteria for identifying differentially-expressed genes from 20E- or DMSO-treated RNA-seq datasets**

Up-regulated genes were defined as those whose expression increased by at least 5-fold and had minimum expression levels of at least 5 Log2 TPM units after 20E treatment. Down-regulated genes were defined as those whose expression reduced by at least 5-fold in the 20E treated condition and had minimum starting expression levels of at least 5 Log2 TPM units in the DMSO control. Functional annotations of *B. mori* gene models were a kind gift from M. Kawamoto, University of Tokyo.

**Criteria for identifying CENP-T enriched or depleted genomic windows**

Genomic windows from 20E-treated or washout datasets with CENP-T level differences that are equal or exceed 3 standard deviations subtracted or added to the mean log2 enrichment score of all bins in the WT were selected. In addition, only those 10 kb windows with total loss or gain of CENP-T from a previously enriched or depleted state, respectively, were considered for our analyses.

**Identification of orthologous genes**

*B. mori* and *T. ni* proteome datasets from Silkbase (http://silkbase.ab.a.u-tokyo.ac.jp, published in (Kawamoto et al., 2019)) and Cabbage Looper Database (https://cabbagelooper.org, published in (Fu et al., 2018) were used. Orthologous genes were selected as those identified as reciprocal best hits in blastp searches (PMID: 2231712) of *B. mori* against *T. ni* proteome and *vice versa*. These analyses revealed 10212 orthologs between the two organisms. We filtered out any orthologs that had multiple hits in the reciprocal organism. This left us with a total of 9533 orthologs for differential expression analyses.

**Criteria for identifying differentially-expressed orthologous genes**

Highly expressed genes for either *B. mori* or *T. ni* were defined as those that had a difference in expression level of at least 5 units to which was added (for *B. mori*) or subtracted (for *T. ni*) the median difference in expression of all 9533 orthologs. Difference in expression was calculated as Log2 TPM+1(*B. mori – T. ni*) for each gene.

**Quantification and statistical analysis**

Statistical details of experiments are detailed in the figure legends. Statistical analyses were performed in RStudio (RStudio Team (2016). RStudio: Integrated Development for R. RStudio, Inc., Boston, MA URL http://www.rstudio.com/).

### Figure S1

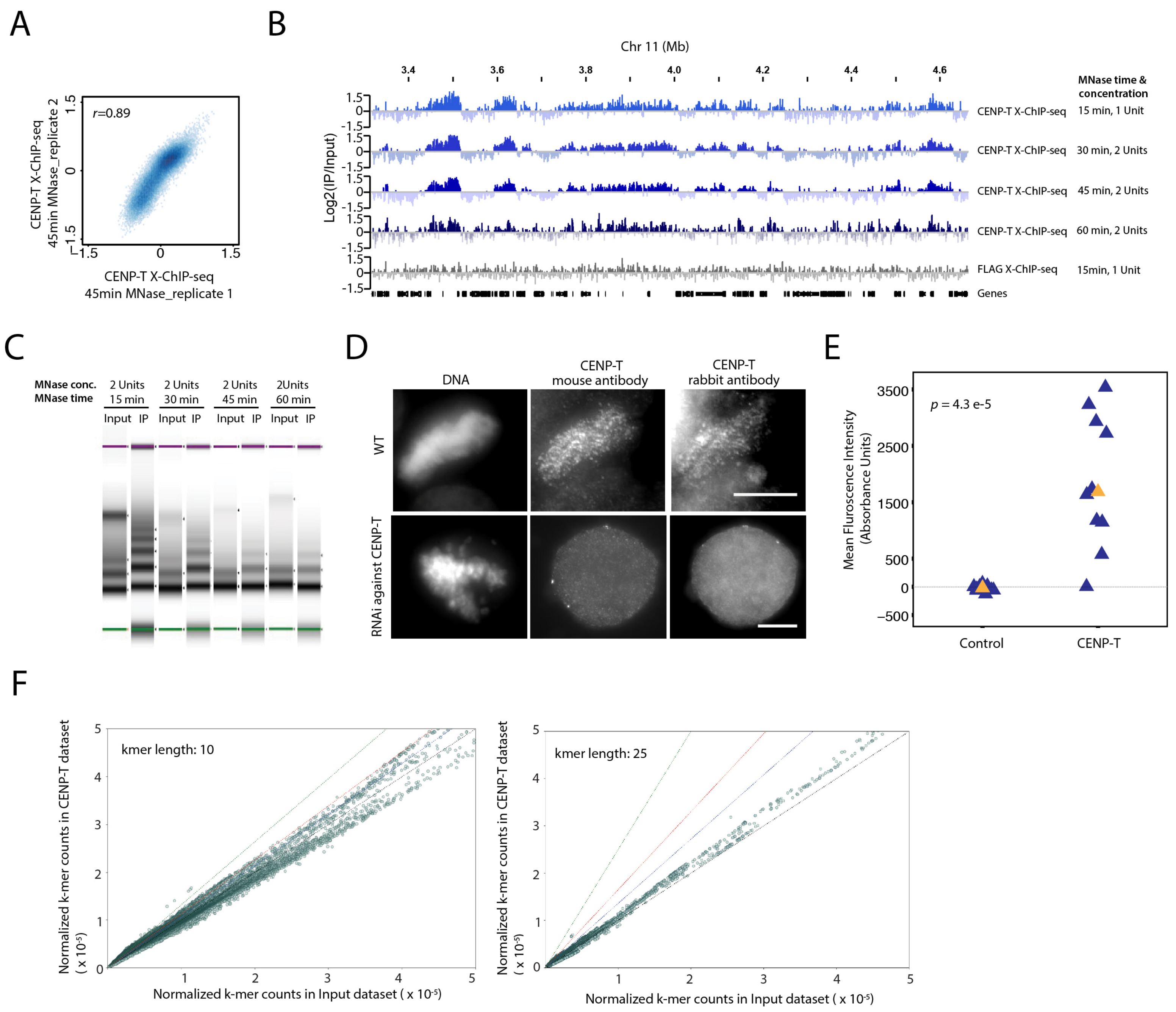

A
B
C
D
E
F
r=0.89
CENP-T X-ChIP-seq 45min MNase_replicate 2
CENP-T X-ChIP-seq 45min MNase_replicate 1
Chr 11 (Mb)
Log2(IP/Input)
MNase time & concentration
CENP-T X-ChIP-seq 15 min, 1 Unit
CENP-T X-ChIP-seq 30 min, 2 Units
CENP-T X-ChIP-seq 45 min, 2 Units
CENP-T X-ChIP-seq 60 min, 2 Units
FLAG X-ChIP-seq 15min, 1 Unit
Genes
MNase conc.
MNase time
2 Units 15 min
2 Units 30 min
2 Units 45 min
2Units 60 min
Input IP
DNA
CENP-T mouse antibody
CENP-T rabbit antibody
WT
RNAi against CENP-T
p = 4.3 e-5
Mean Fluroscence Intensity (Absorbance Units)
Control
CENP-T
kmer length: 10
kmer length: 25
Normalized k-mer counts in CENP-T dataset ( x 10-5)
Normalized k-mer counts in Input dataset ( x 10-5)

### Figure S2

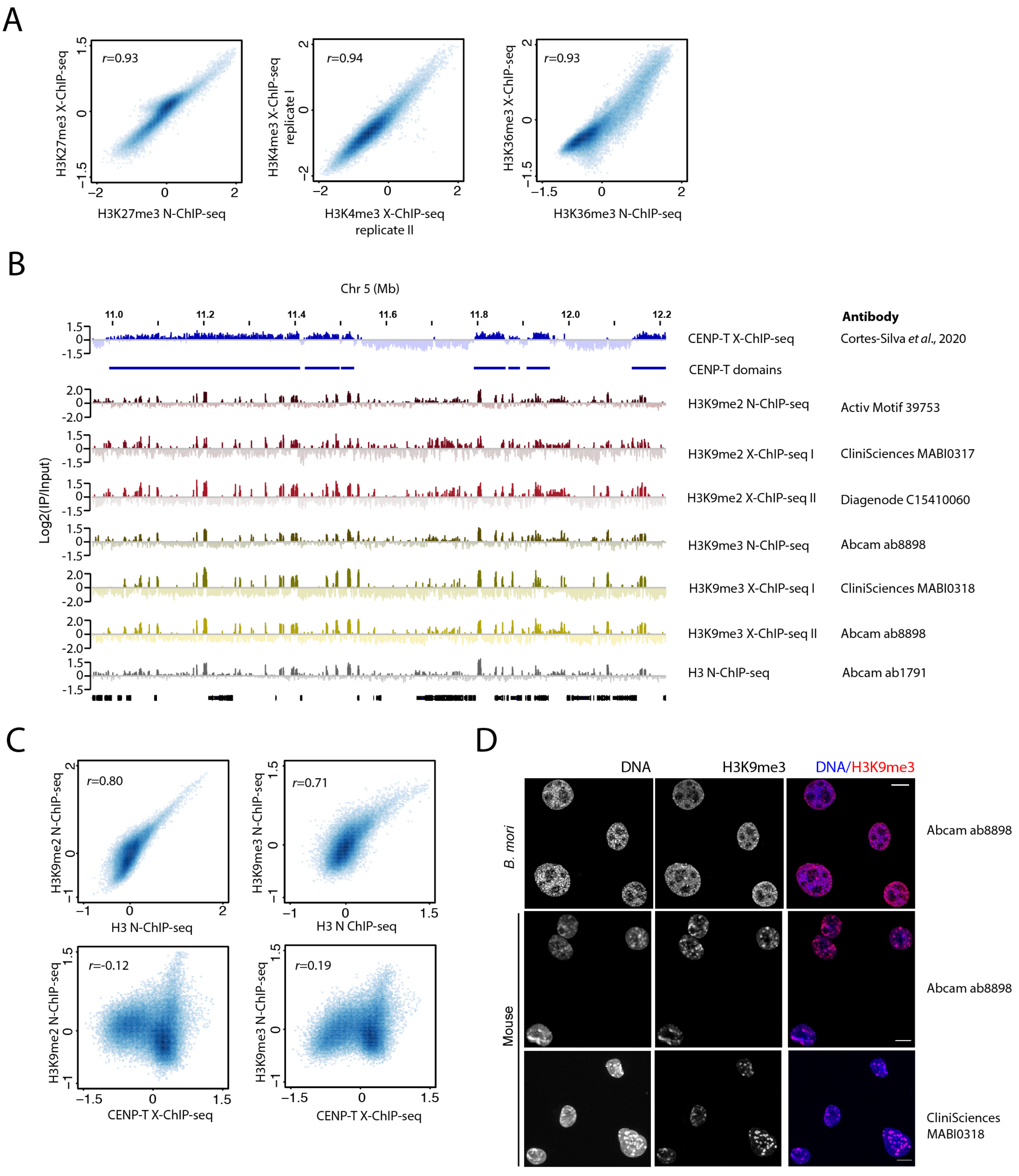
A
r=0.93
H3K27me3 X-ChIP-seq
H3K27me3 N-ChIP-seq
r=0.94
H3K4me3 X-ChIP-seq replicate I
H3K4me3 X-ChIP-seq replicate II
r=0.93
H3K36me3 X-ChIP-seq
H3K36me3 N-ChIP-seq
B
Chr 5 (Mb)
11.0
11.2
11.4
11.6
11.8
12.0
12.2
Antibody
Log2(IP/Input)
CENP-T X-ChIP-seq
Cortes-Silva et al., 2020
CENP-T domains
H3K9me2 N-ChIP-seq
Activ Motif 39753
H3K9me2 X-ChIP-seq I
CliniSciences MABI0317
H3K9me2 X-ChIP-seq II
Diagenode C15410060
H3K9me3 N-ChIP-seq
Abcam ab8898
H3K9me3 X-ChIP-seq I
CliniSciences MABI0318
H3K9me3 X-ChIP-seq II
Abcam ab8898
H3 N-ChIP-seq
Abcam ab1791
C
r=0.80
H3K9me2 N-ChIP-seq
H3 N-ChIP-seq
r=0.71
H3K9me3 N-ChIP-seq
H3 N ChIP-seq
r=-0.12
H3K9me2 N-ChIP-seq
CENP-T X-ChIP-seq
r=0.19
H3K9me3 N-ChIP-seq
CENP-T X-ChIP-seq
D
DNA
H3K9me3
DNA/H3K9me3
B. mori
Mouse
Abcam ab8898
Abcam ab8898
CliniSciences MABI0318

### Figure S3

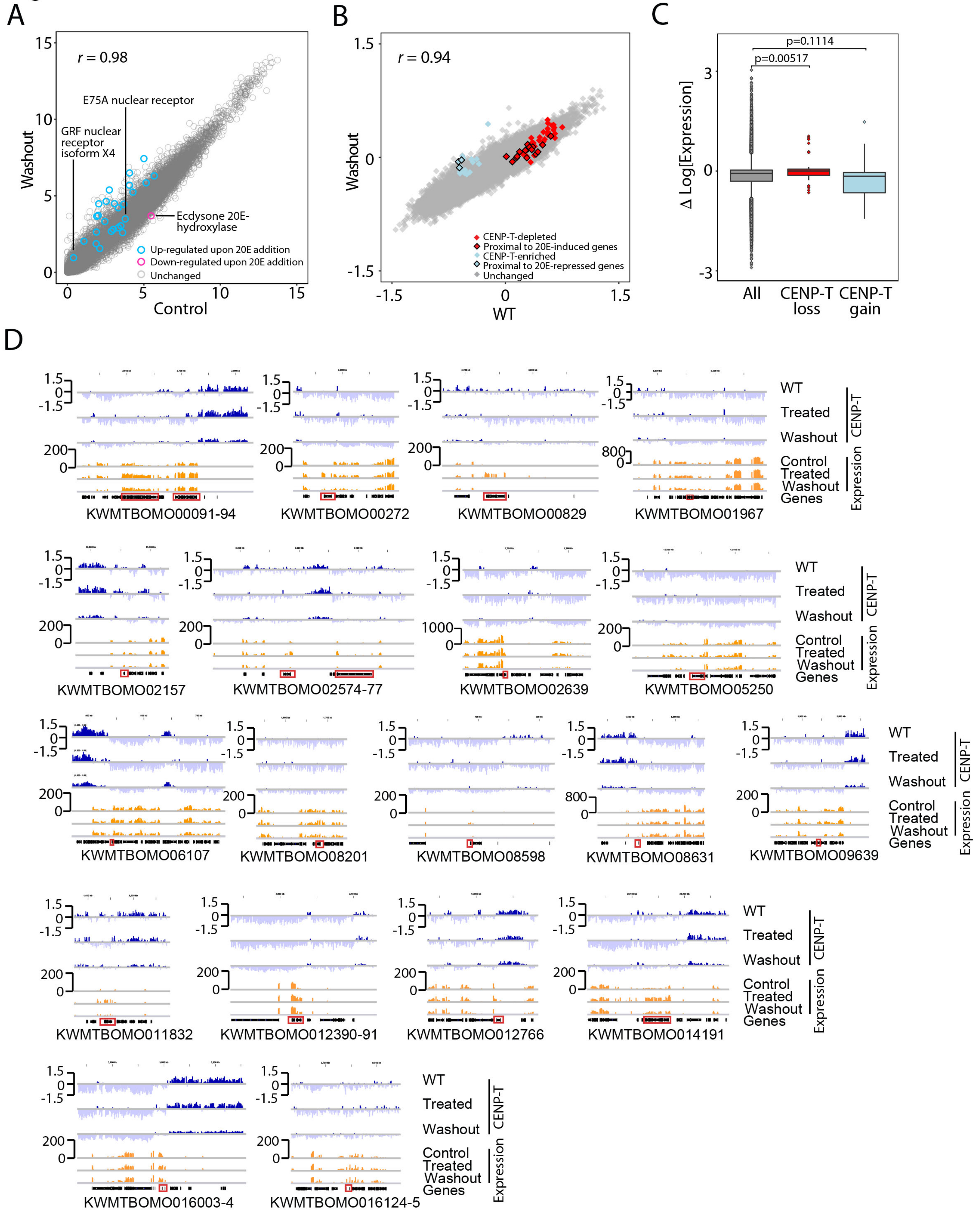
A
r = 0.98
Washout
Control
GRF nuclear receptor isoform X4
E75A nuclear receptor
Ecdysone 20E-hydroxylase
Up-regulated upon 20E addition
Down-regulated upon 20E addition
Unchanged
B
r = 0.94
Washout
WT
CENP-T-depleted
Proximal to 20E-induced genes
CENP-T-enriched
Proximal to 20E-repressed genes
Unchanged
C
Δ Log[Expression]
p=0.1114
p=0.00517
All
CENP-T loss
CENP-T gain
D
WT
Treated
Washout
CENP-T
Control
Treated
Washout
Genes
Expression
KWMTBOMO00091-94
KWMTBOMO00272
KWMTBOMO00829
KWMTBOMO01967
KWMTBOMO02157
KWMTBOMO02574-77
KWMTBOMO02639
KWMTBOMO05250
KWMTBOMO06107
KWMTBOMO08201
KWMTBOMO08598
KWMTBOMO08631
KWMTBOMO09639
KWMTBOMO011832
KWMTBOMO012390-91
KWMTBOMO012766
KWMTBOMO014191
KWMTBOMO016003-4
KWMTBOMO016124-5

### Figure S4

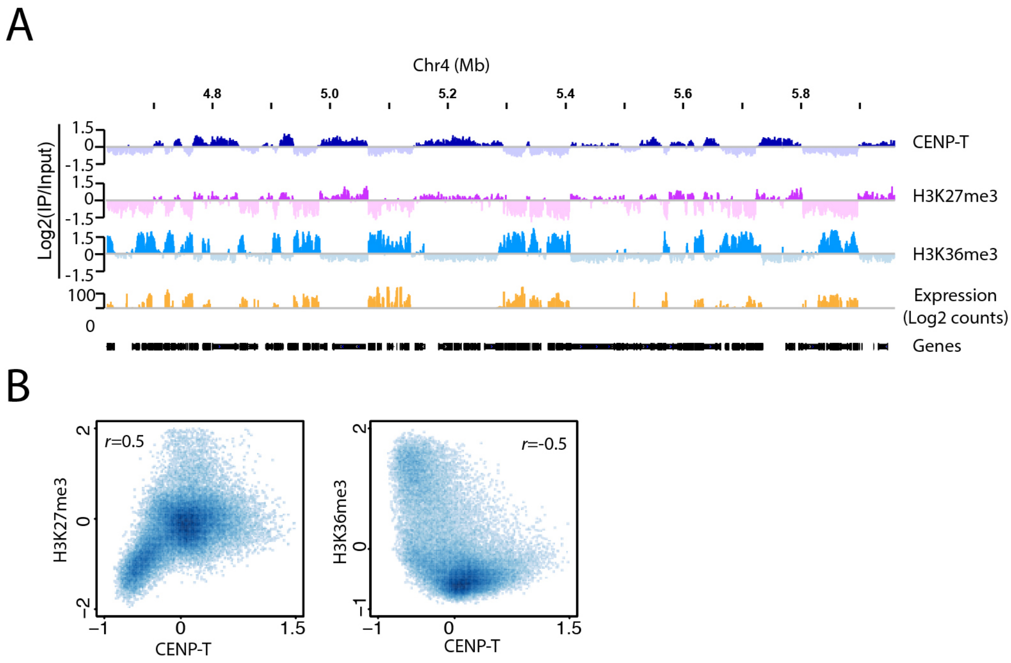
A
Chr4 (Mb)
4.8
5.0
5.2
5.4
5.6
5.8
Log2(IP/Input)
1.5
0
-1.5
CENP-T
H3K27me3
H3K36me3
100
0
Expression
(Log2 counts)
Genes
B
r=0.5
H3K27me3
CENP-T
r=-0.5
H3K36me3
CENP-T
